# Structure of a heteropolymeric type 4 pilus from a monoderm bacterium

**DOI:** 10.1101/2023.06.15.545089

**Authors:** Robin Anger, Laetitia Pieulle, Meriam Shahin, Odile Valette, Hugo Le Guenno, Artemis Kosta, Vladimir Pelicic, Rémi Fronzes

## Abstract

Type 4 pili (T4P) are important virulence factors, which belong to a superfamily of nanomachines ubiquitous in prokaryotes, called type 4 filaments (T4F). T4F are defined as helical polymers of type 4 pilins. Recent advances in cryo-electron microscopy (cryo-EM) led to structures of several T4F. This revealed that the long N-terminal α-helix, the trademark of pilins, packs in the centre of the filaments to form a hydrophobic core, which in bacteria is accompanied by the melting (unfolding) of a portion of α1. Since all available bacterial T4F structures are from diderm species, we tested whether this architecture is conserved in phylogenetically distant species by determining the structure of the T4P of the monoderm *Streptococcus sanguinis*. Our 3.7 A resolution cryo-EM structure of this heteropolymeric T4P, and the resulting full atomic model including all minor pilins, highlight universal features of bacterial T4F and have widespread implications in understanding their biology.

## Introduction

Type 4 filaments (T4F) are a superfamily of filamentous nanomachines ubiquitous in Bacteria and Archaea^1, 2^. The best known T4F are type 4 pili (T4P)^3^ and type 2 secretion systems (T2SS)^4^. T4F mediate a staggering array of different functions such as adhesion, motility (swimming and twitching), DNA uptake, formation of bacterial communities, and protein secretion^1^. T4F have been an important research area for decades because they are virulence factors in many bacterial pathogens^3^.

T4F are filamentous polymers of type 4 pilins, usually composed of one major and several minor (low abundance) subunits^5^. These subunits are synthesised with a distinctive N-terminal (NT) class 3 signal peptide (SP3)^6^, which needs to be processed by a dedicated prepilin peptidase before T4F assembly^7, 8^. The SP3 consists of a hydrophilic leader peptide, followed by a stretch of 20-25 predominantly hydrophobic residues^5^ with often a Glu in position 5. In the characteristic “lollipop” structure of pilins, this hydrophobic stretch represents the “stick” that protrudes from a globular head, which is usually centred on an anti-parallel β-sheet^9^. The stick is the NT half of an α1-helix of 50-55 residues (α1N), the C-terminal (CT) half of which (α1C) is packed within the globular head. The first atomic model of a T4P *in Neisseria gonorrhoeae*^9^, based on fibre diffraction and electron microscopy results, suggested that the α1 helices form a hydrophobic core by packing helically, in a roughly parallel fashion. This model was then refined to 12 Å resolution by fitting the crystal structure of the pilin into a cryo-electron microscopy (cryo-EM) density map^10^. In the past few years, advances in cryo-EM led to multiple near-atomic resolution T4F structures^11^, which revealed a conserved helical architecture, albeit with different symmetry parameters. A striking observation in *N. meningitidis* T4P was that a portion of α1N is “melted” (non-helical), which must occur during polymerisation of the pilin subunits into filaments^12^. This feature was subsequently reported in T4P from enterohemorrhagic *Escherichia coli*^13^ (EHEC), *Pseudomonas aeruginosa*^14^, *N. gonorrhoeae*^14^ and *Thermus thermophilus*^15^, and in T2SS from *Klebsiella oxytoca*^16^ and *Geobacter sulfurreducens*^17^.

All the above structures were determined in diderm bacterial species, which were for more than 20 years the only available models to study T4F. It therefore remains to be determined whether this filament architecture is universal in bacteria. Moreover, none of these structures provided a complete picture of the corresponding T4F since they do not encompass minor pilins, which are key players in T4F biology, often poorly characterised. Recently, phylogenetically distant monoderm bacteria became a promising new T4F research avenue^18, 19^, which could help answering the above issues. The cutting-edge monoderm model is *Streptococcus sanguinis*, a commensal of the human oral cavity frequently causing endocarditis. *S. sanguinis* T4P have been characterised in depth^20–24^, revealing that they are heteropolymers of two major pilins (PilE1, PilE2) – which is unusual – and three minor pilins (PilA, PilB, PilC). The structure and function of all these subunits has been determined, which is yet to be achieved for most T4F. The two major subunits display a canonical pilin fold, with an uncommon highly flexible C-terminus^22^. The three minor pilins are predicted to form a complex that localises at the T4P tip^24^ and promotes adhesion to various host receptors via the modular pilins PilB and PilC^23, 24^. PilB and PilC are unusually large pilins with grafted modules conferring adhesive properties^23, 24^, while PilA is an anchor for PilC at the tip of pilus, *S. sanguinis* T4P thus look particularly suitable for determining whether the filament architecture with a melted portion of α1N is universal in bacterial T4F, and for producing a complete picture of a T4F including all its subunits. This was achieved here by using cryo-EM and modelling, and we discuss the wider implications of our findings for this 3 superfamily of nanomachines.

## Results

### T4P on S. sanguinis are flexible filaments 7 nm in width

We previously reported that *S. sanguinis* T4P in highly pure pilus preparations exhibit two different morphologies: thick/wavy (12-15 nm) and thin/straight (6-7 nm)^20, 22^. To determine the morphology of T4P on the surface of bacterial cells, we observed *S. sanguinis* cells by transmission EM (TEM) after negative staining. To facilitate observation of T4P, we used a Δ*fim* mutant where we deleted the *fim* locus, which is involved in the production of the unrelated sortase-assembled pilus^25^. As can be seen in Fig. 1, T4P emanating from *S. sanguinis* cells exhibit a classical T4P morphology^1^. They are (1) predominantly polar, (2) up to several µm in length, and (3) 6.78 ± 0.31 nm in diameter. Occasionally, a couple of filaments aggregate laterally (Fig. 1), but they do not form large bundles like T4P in most diderm species^1^. Therefore, in pilus preparations, the 6-7 nm-wide filaments correspond to native *S. sanguinis* T4P.

**Fig. 1.**
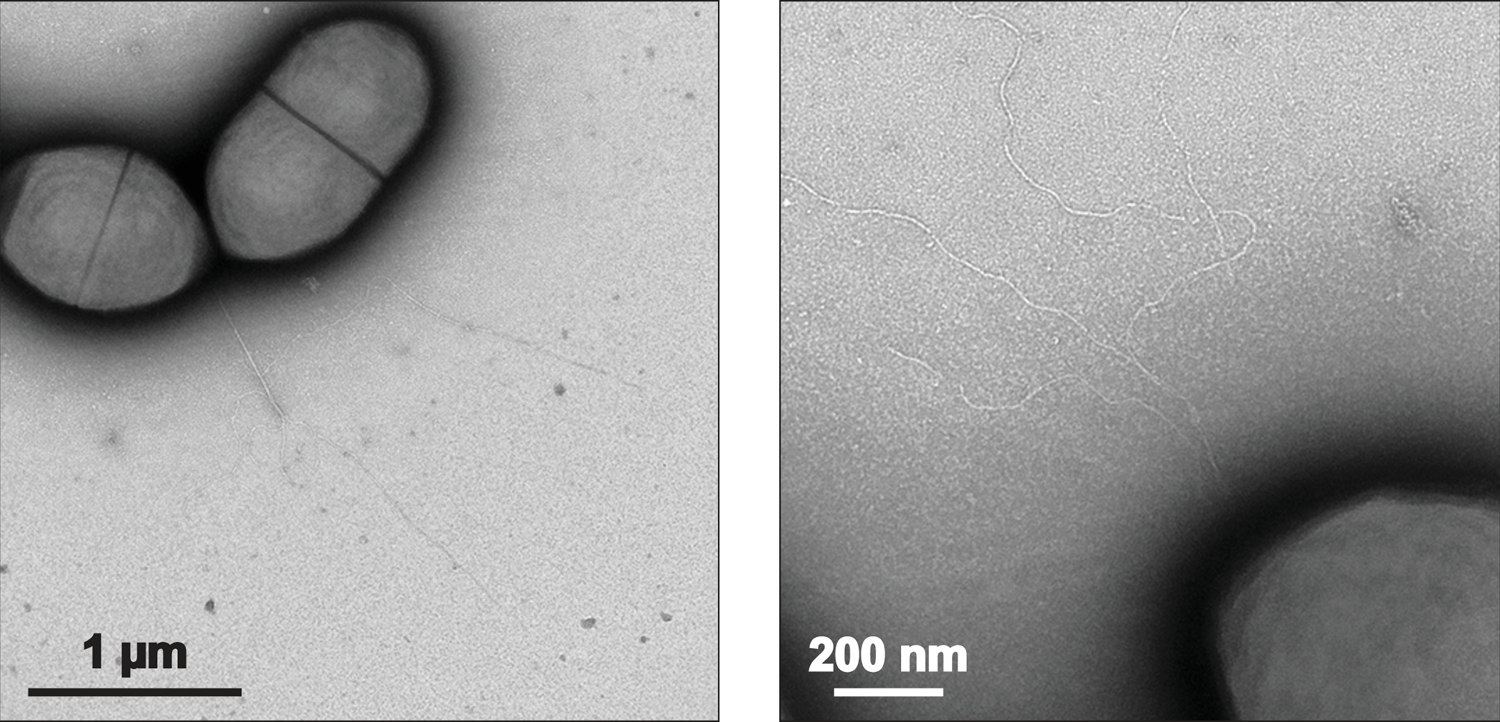
Imaging T4P on *S. sanguinis* cells by TEM after negative staining. Two representative images, at different scales, are shown.

To determine the structure of *S. sanguinis* T4P, we analysed purified pili by cryo-EM. Three distinct types of filaments were observed on the micrographs: the two different forms of T4P previously seen by TEM^20, 22^ – 6-7 nm and 12-15 nm in diameter – as well as very thin filaments (3 nm-wide) (Fig. S1A). A preliminary structural analysis of the latter two filaments allowed us to exclude them from further analysis. In brief, for the 12-15 nm-wide filaments, we generated a low-resolution electron density map after 2D classification, which revealed a cylindrical structure with no distinctive features (Fig. S1B). It remains thus unknown whether these thick T4P result from a dramatic change in quaternary conformation of the filaments (similarly to what has been reported for *N. gonorrhoeae* T4P^26^) or from their limited denaturation upon shearing from the bacterial surface. In contrast, for the 3 nm-wide filaments, we could produce a density map at 6 Å resolution in which the structure of B-DNA could be fitted readily (Fig. S1C), indicating that these thin filaments correspond to DNA. This extracellular DNA, which is unlikely to result from cell lysis – because no ribosomes were observed in the pilus preparations – is probably actively released by *S. sanguinis*, a well-known property of this species^27^.

We then focused on the 6-7 nm-wide filaments, which look like the T4P seen on the surface of *S. sanguinis*. We segmented these filaments and performed iterative rounds of 2D classification followed by *ab initio* reconstruction. After 3D refinement and local refinement on the central portion along the filament axis (150 Å long), we generated a final density map at 3.7 Å resolution, as estimated by gold-standard Fourier shell correlation (FSC) (Fig. 2A). The resolution could not be improved by additional 3D classification/variability experiments. The reconstructed filament is a cylinder, 70 Å in diameter, with significant curvature (Fig. 2B). The resolution, which is close to 3.3 Å in the central portion of the filament, decreases to 4.3 Å towards the edges, especially towards both ends (Fig. 2C). This makes it impossible to precisely calculate the helical symmetry operators for the filament, which could nevertheless be estimated at 11 Å in rise and 93° in twist angle. The 3D reconstructions were not improved further upon imposition of these helical symmetry operators (Fig. S2).

**Fig. 2.**
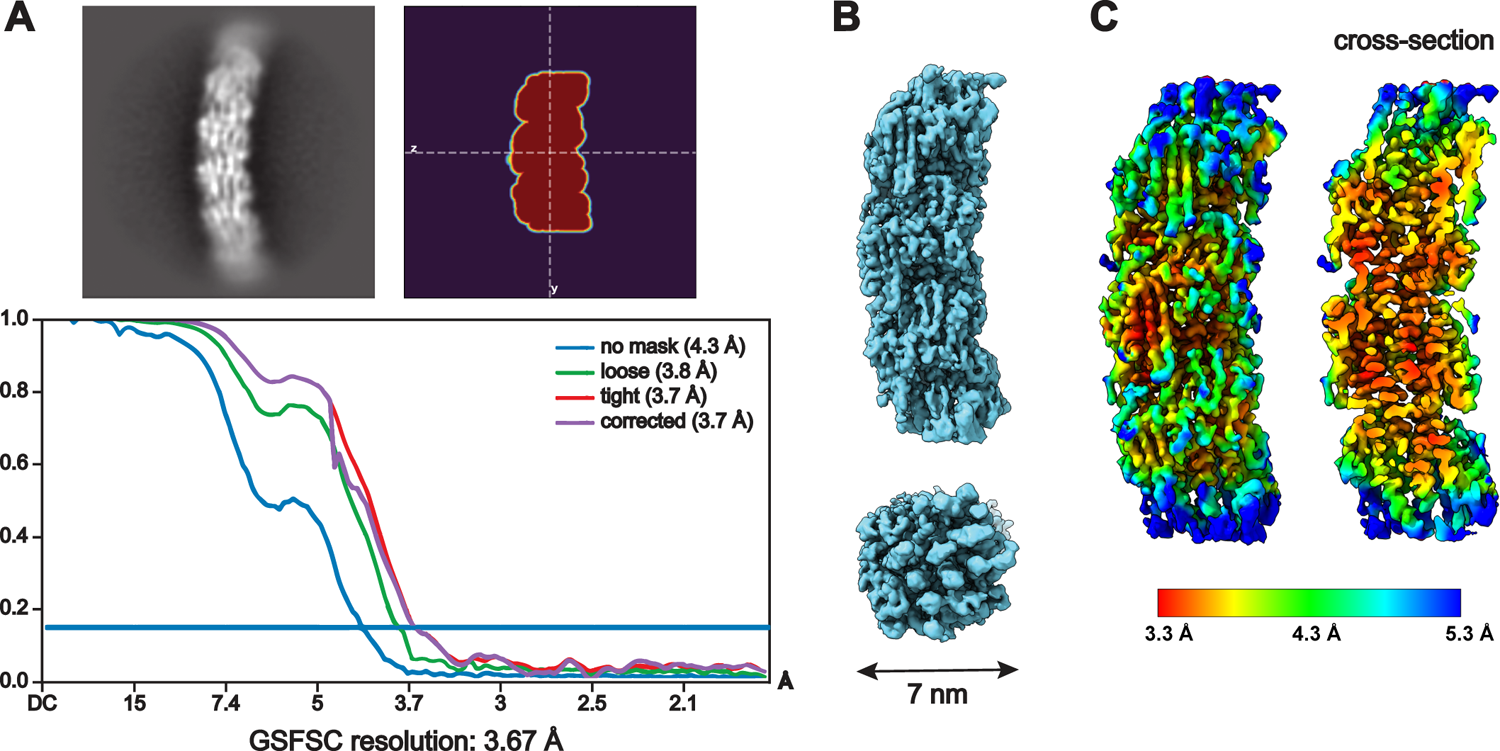
Cryo-EM density map of *S. sanguinis* T4P at 3.7 Å resolution. The map was obtained without imposing a symmetry. **A**) **Left**, representative average of pilus sections from cryo-EM micrographs after 2D classification. **Right**, slice of the mask used for 3D refinement of the central portion. **Bottom**, estimation of the resolution by gold standard FSC. **B**) Final density map sharpened with DeepEMhancer^35^. Side and end views of the filament are presented, with its diameter indicated. **C**) Final density map coloured according to local resolution. A cross-section is shown on the right.

### S. sanguinis T4P are composed of two major subunits arranged stochastically

In contrast to the other characterised bacterial T4F, *S. sanguinis* T4P are heteropolymers composed of two major pilins in comparable amounts, PilE1 and PilE2^20, 22^. These two subunits show extensive sequence identity (Fig. 3A). The main distinctive feature is an extra 8-aa loop at the end of the αβ region in PilE1, which makes this protein slightly longer than PilE2. Within the final density map, individual pilin subunits could be readily identified (Fig. 3B). Strikingly, the regions of difference between PilE1 and PilE2 – a CT “tail” and the 8-aa loop between residues 92 and 100 in PilE1 – correspond to areas of significantly lower quality in the density map (Fig. S3). At the resolution of the map, it should have been possible to model the 8-aa loop, which in contrast could not be modelled at all. The possibility that this might be due to an inherent flexibility of the 8-aa loop can be excluded by the previously determined NMR structure of PilE1, which showed no flexibility in this portion of the structure within the NMR ensemble^22^. Rather, the lower quality in the density map likely results from the stochastic assembly (not in a specific order) of PilE1 and PilE2 within *S. sanguinis* T4P.

**Fig. 3.**
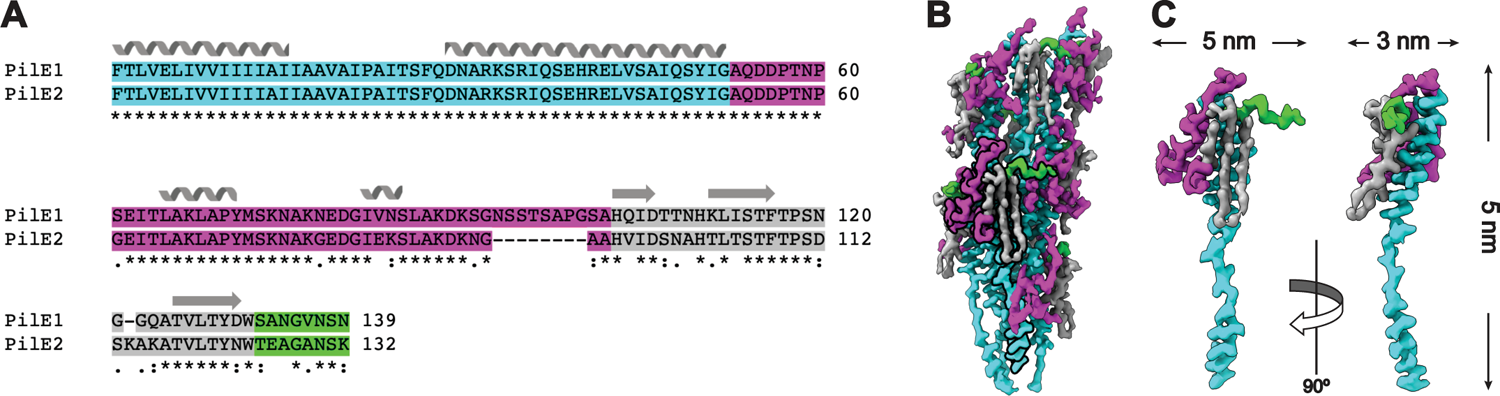
*S. sanguinis* T4P are canonical T4F. **A**) Sequence alignment between PilE1 and PilE2 using ClustalOmega. (*) conserved aa; (:) aa with strongly similar properties; (.) aa with weakly similar properties. The NT α1-helix, αß-loop, antiparallel ß-sheet, and CT tail are coloured in cyan, magenta, grey and green, respectively. The secondary structures are indicated above the sequences. **B**) Pilins can be readily visualised in the density map. The same colour code is used as in panel A. Densities corresponding to a pilin are underlined with bold contours. **C**) Individual pilin shown in orthogonal orientations. The protruding portion of the α1 helix is clearly interrupted by an unfolded stretch. The same colour code is used as in panel B.

Recently, the structure of a heteropolymeric T4F has been reported – the archaellum (archaeal flagellum) from *Methanocaldococcus villosus* – in which the two major subunits ArlB1 and ArlB2 alternate regularly^28^. Such filament assembly was proposed to result from the polymerisation of pre-formed ArlB1ArlB2 heterodimers, since the two proteins interact preferentially with one another^28^. We therefore tested the interactions between *S. sanguinis* PilE1 and PilE2 using the bacterial adenylate cyclase two-hybrid (BACTH) system^29^, which has proven effective for many T4F proteins^30, 31^. For each pilin, we generated two different BACTH plasmids by fusing full-length PilE1 and PilE2 at the NT of T18 and T25 domains of *Bordetella pertussis* adenylate cyclase. We then assessed functional complementation between all possible pairs of T18 and T25 plasmids by co-transformation in an *E. coli cya* mutant and plating on selective indicator plates. As can be seen in Fig. 4A, all plasmid combinations yielded coloured colonies, indicating that PilE1 and PilE2 interact. The efficiency of the functional complementation, which reflects the strength of the protein-protein interaction^29, 30^, was quantified by measuring β-galactosidase activities (Fig. 4B). In contrast with what was reported for ArlB1 and ArlB2 in *M. villosus*^28^, we found that PilE1 and PilE2 interact equally well with themselves as with one another. This argues against the possibility that *S. sanguinis* T4P could be heteropolymers in which the two major subunits would alternate regularly upon polymerisation of pre-formed PilE1PilE2 heterodimers. Rather, our findings suggest that the major subunits PilE1 and PilE2 are distributed stochastically in the filaments.

**Fig. 4.**
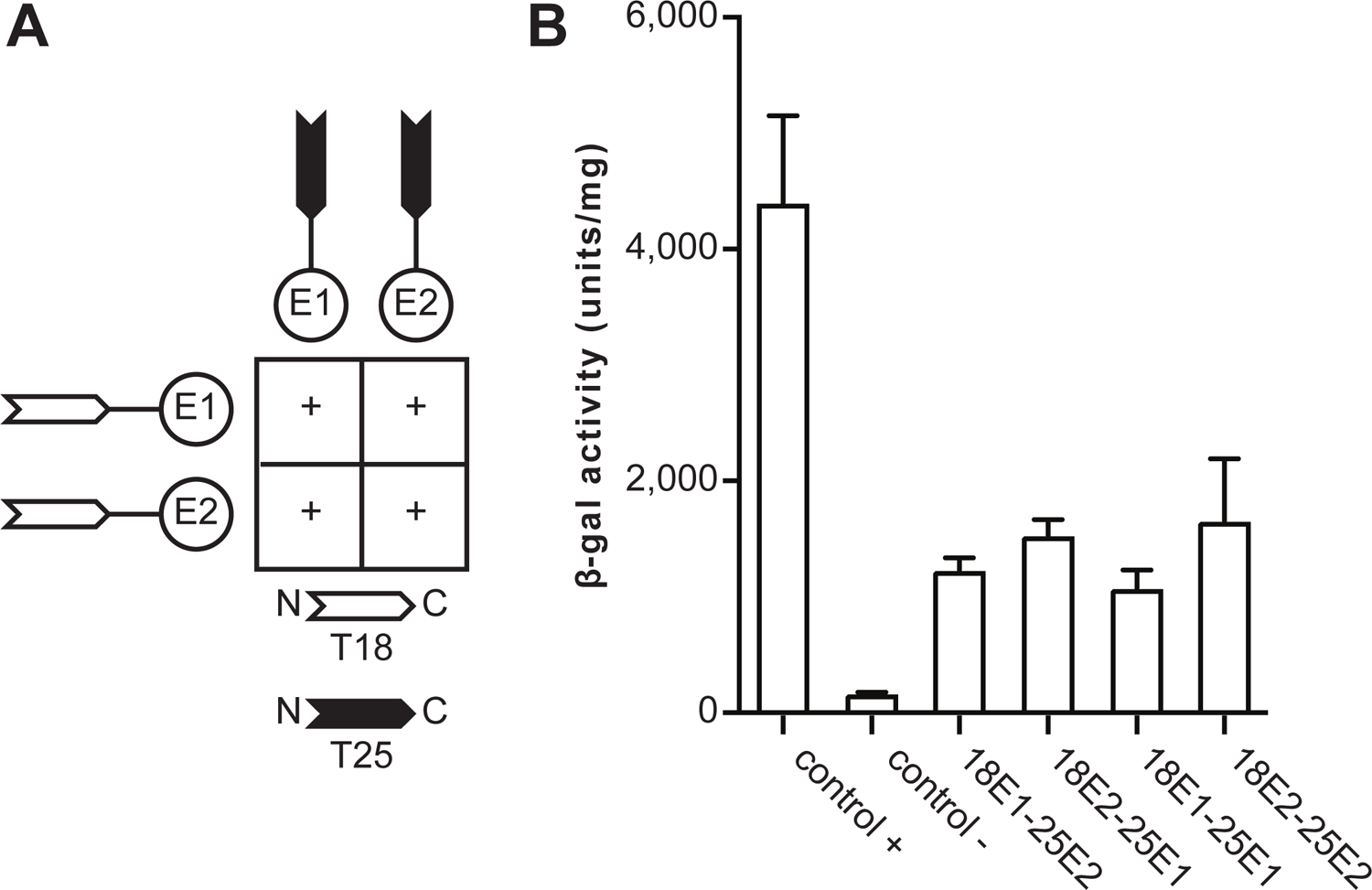
Testing the interactions between PilE1 and PilE2 using BACTH. The T18 and T25 domains of *B. pertussis* adenylate cyclase were fused to the N-terminus of the full-length proteins. T18 and T25 plasmids pairs were co-transformed in an *E. coli cya* mutant, before plating on selective indicator plates. **A**) Combinations of T18 and T25 plasmids producing coloured colonies (+), which indicates an interaction between the corresponding PilE1 or PilE2 proteins. **B**) The strength of the interactions was quantified by measuring β-galactosidase activities (U/mg). The results are the average ± SD from three independent experiments.

### S. sanguinis T4P are canonical bacterial T4F

The resolution of the map was better than 3.4 Å in most of the parts, allowing an accurate building of the filament. The resolution was highest in the central portion of the filament, which corresponds to the α1 helices. This allowed us to fit readily the peptide backbone of that portion of the pilin – identical in sequence in PilE1 and PilE2 (Fig. 3A) – within the representative cryo-EM densities (Fig. 5A). Fitting was unambiguous and large sidechains were readily visible. Critically, in each subunit the α1 helix is interrupted by an unfolded stretch (Fig. 5A), showing that this feature is conserved in all bacterial T4F structures determined so far^11^.

**Fig. 5.**
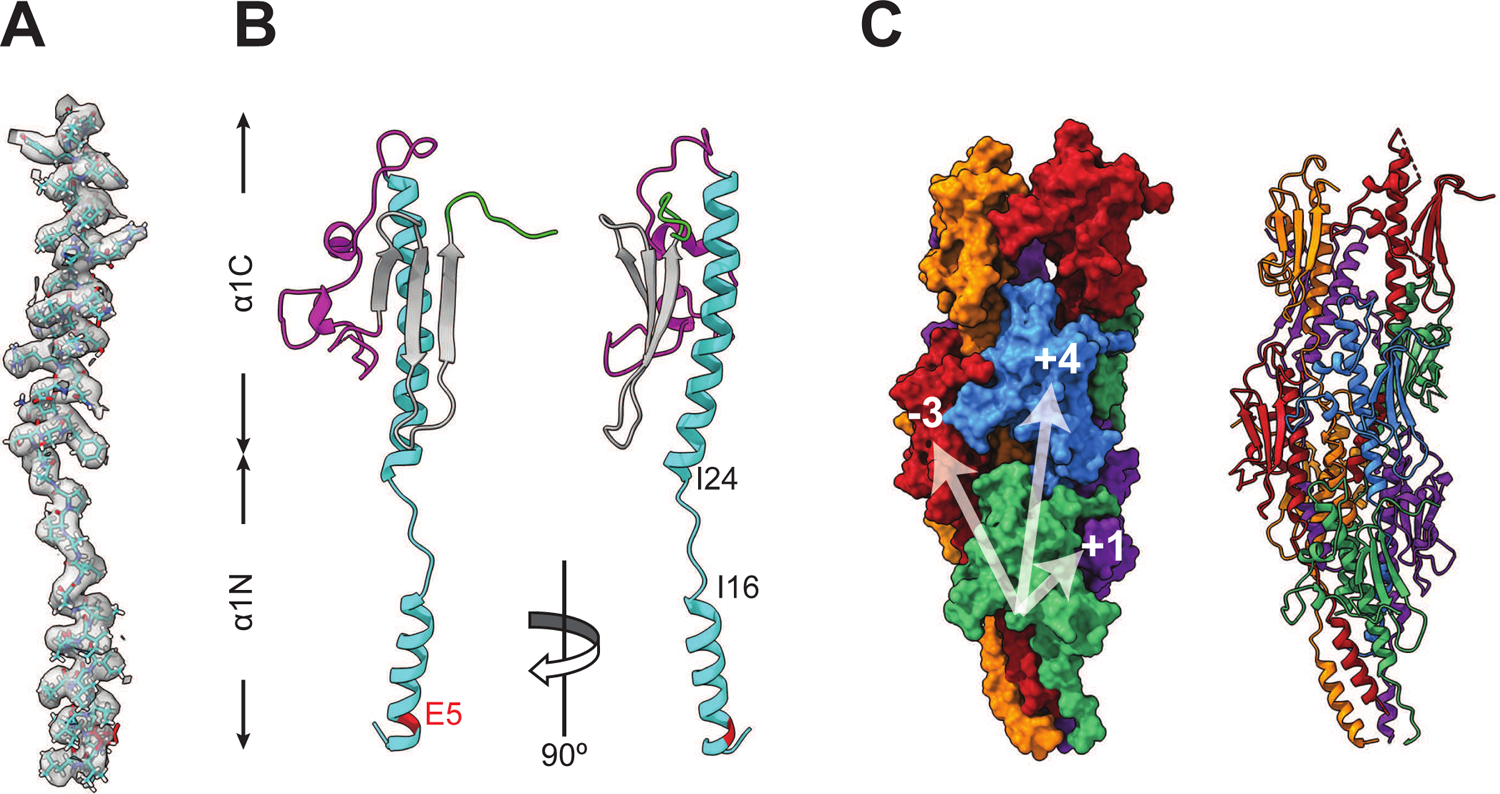
Atomic model of *S. sanguinis* T4P. **A**) Fitting of PilE1/PilE2 identical NT portion (stick representation) within the representative cryo-EM densities in the final density map. **B**) Ribbon representation of a pilin subunits in the filament. The positions of Glu_5_ and the melted region between Ile_16_ and Ile_24_ are indicated. We used the same colour code as in Fig. 3. **C**) *S. sanguinis* T4P structure. Surface (left) and ribbon (right) representations with the individual pilins outlined in different colours. Connectivity is shown in the right-handed 1-start (+1), right-handed 4-start (+4), and left-handed 3-start (−3) helices.

As PilE1 and PilE2 share high sequence identity, we arbitrarily chose PilE1 to produce an atomic model of *S. sanguinis* T4P. Copies of PilE1 generated by AlphaFold^32^ were manually docked into the final density map. Flexible fitting of each subunit and optimisation of the model geometry was performed using ISOLDE^33^ in ChimeraX^34^, against a sharpened map obtained with DeepEMhancer^35^. The final model was refined in Phenix^36^ against a map automatically sharpened using the same program. In our atomic model, the pilin subunits display a canonical lollipop structure in which the α1N portion of a 52 residues-long α1 helix protrudes from a globular head (Fig. 5B). In the globular head, α1C is packed against a three-strand β-sheet, which is flanked at its N- and C-termini by regions with no major distinctive secondary structure elements, *i.e.*, a large αβ-loop and a short CT tail, respectively (Fig. 5B). Critically, as in other structures of bacterial T4F^11^, a portion of α1N is melted (Fig. 5B), the extent of which varies slightly between Ile_16_ and Gln_28_ in the different chains. The superposition of the structures of the complete pilins that we were able to resolve, shows that the portion of α1N up to the end of the melted region is intrinsically flexible (Fig. S4). In contrast, the other parts of the pilin remain almost unchanged, aligning with 1.3 Å root-mean-square deviation (RMSD) globally.

The overall architecture of *S. sanguinis* T4P conforms to the shared structural principles in T4F. In brief, the hydrophobic core of the filament consists of a bundle of α1 helices, leaving the opposite face of the globular heads to form the outer shell (Fig. 5C). The filaments are right-handed with four subunits-turn, which display approx. 11 Å rise and 93° twist between consecutive subunits within the 1-start helix (Fig. 5C). The interactions between subunits that hold the filament together involve residues in various parts of the pilin (Fig. S5) and can be recapitulated at the levels of the 1-start, 3-start and 4-start helices (Fig. 6). Perhaps the most unusual interaction is within the right-handed 1-start helix, where the CT tail in PilE1 (mostly negatively charged) engages the next pilin αβ-loop (mostly positively charged) through electrostatic interactions (Fig. 6A). This stabilising interaction therefore resembles a “velcro” closure mechanism. In contrast, in the structure of PilE1 monomers previously determined by NMR^22^, this CT tail is highly flexible. At the level of the 1-start helix, there are also hydrophobic contacts between the α1N helices of the S_-1_, S and S_+1_ subunits. Phe1, Leu_6_ and Val_9_ of subunit S thus interact with Phe_1_ of S_+1_, and Ile_12_ and Ile_13_ of S_-1_ (Fig. 6A).

**Fig. 6.**
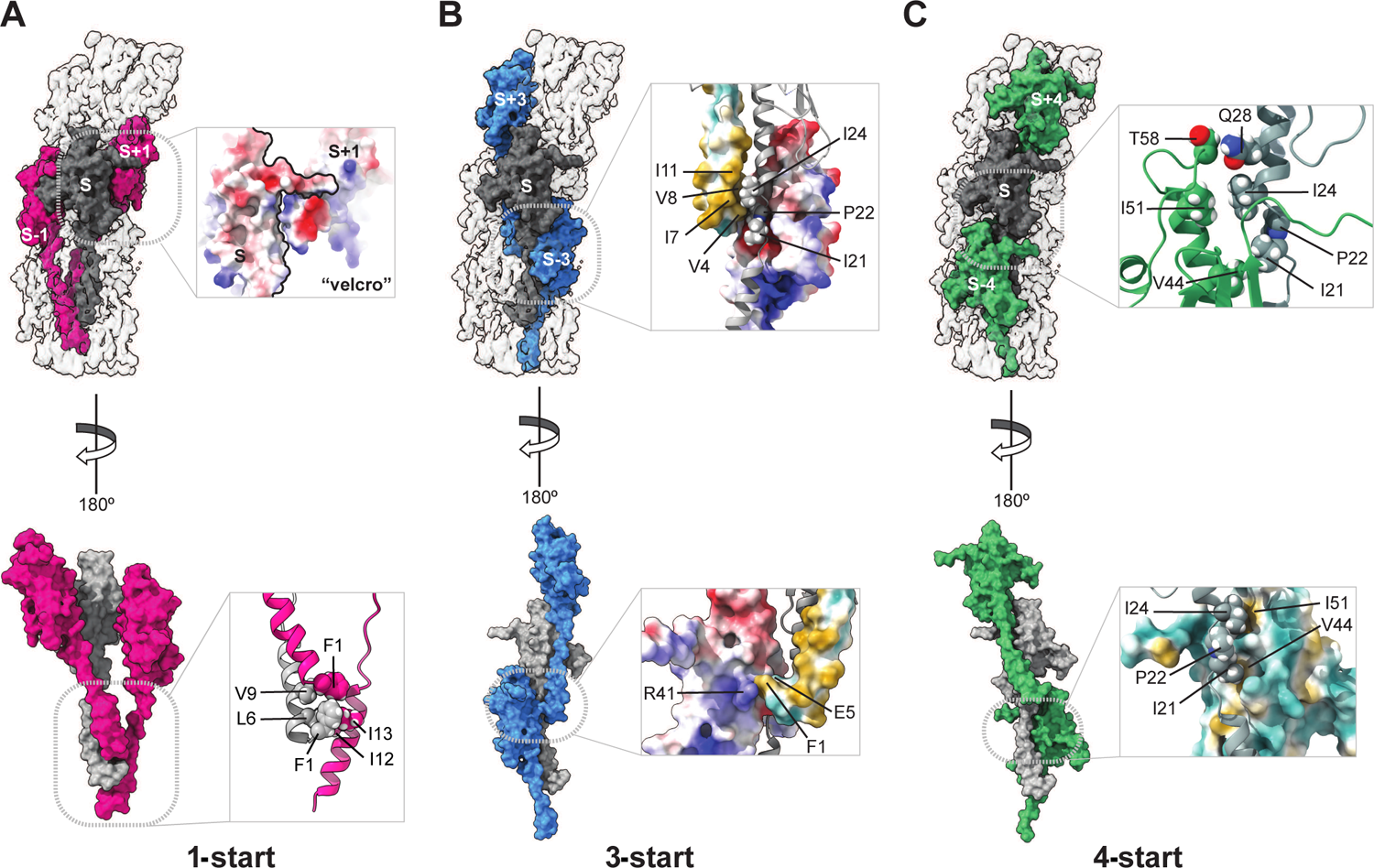
Interactions between the subunits in *S. sanguinis* T4P. **A**) Interactions between the S (grey) and S_+1_/S_-1_ subunits (pink) in the 1-start helix (180° views). **Top**, enlarged view shows a surface representation of the electrostatic potential of the S and S_+1_ subunits that interact via a velcro mechanism. **Bottom**, enlarged view of the interactions between the NT parts of the three pilins (ribbon representation), with the residues involved highlighted. **B**) Interactions between the S (grey) and S_+3_/S_-3_ subunits (blue) in the 3-start helix (180° views). **Top**, enlarged view of the interactions between the S (ribbon representation), S_+3_ (hydrophobicity surface representation) and S_-3_ (electrostatic potential surface representation) subunits, with the residues involved highlighted. **Bottom**, enlarged view of the interactions between the S_+3_ (hydrophobicity surface representation) and S_-3_ (electrostatic potential surface representation) subunits, with the residues involved highlighted. **C**) Interactions between the S (grey) and S_+4_/S_-4_ subunits (green) in the 4-start helix (180° views). **Top**, enlarged view of the interactions between the S and S_-4_ subunits (both in ribbon representation), with the residues involved highlighted. **Bottom**, enlarged view of the interactions between the S (ribbon representation) and S_-4_ (hydrophobicity surface representation) subunits is represented, and the interacting residues are indicated.

At the level of the left-handed 3-start, the melted region in subunit S is held in a groove between α1N of subunit S_+3_, and α1C of subunit S_-3_ (Fig. 6B). The Ile_21_ and Ile_24_ residues of subunit S establish hydrophobic contacts contact with Val_4_, Ile_7_, Val_8_ and Ile_11_ of subunit S_+3_, helping to break the helix symmetry around Pro_22_ (Fig. 6B). The hydrophobic interactions between S and S_+3_ is favoured by the other side of the groove, the back of α1C of S_-3_, which is mostly negatively charged (Fig. 6B). Phe_1_ of subunit S_+3_ is thus held between charged residues Arg_41_ of S_-3_, and Glu_5_ of S_+3_ (Fig.6B). Critically, Glu_5_ – whose essential role in T4F assembly^5^ remains incompletely understood^37^ – defines together with Arg_41_ (and Thr_2_), a charged path at the centre of the pilus core. The electrostatic interactions between these three residues are strengthened by the hydrophobic shell that surrounds them. Finally, in the 4-start helix, the Ile_21_ and Ile_24_ residues in subunit S also establish hydrophobic interactions with Val_44_ and Ile_51_ in S_-4_ (Fig. 6C). In addition, Gln_28_ in subunit S establishes electrostatic interactions with Thr_58_ at the top of the α1 helix in subunit S_-4_ (Fig. 6C).

Taken together, these findings indicate that *S. sanguinis* T4P are canonical bacterial T4F, *i.e*., the centre of the filament is formed by the packing of α1 helices, a portion of which is unfolded. Since these structural features – previously observed in T4F in diderm species – are conserved in a distant monoderm species such as *S. sanguinis*, they are likely to be universal in Bacteria.

#### Full structural model of S. sanguinis T4P encompassing the minor pilins

We previously determined the structure and function of the three other subunits of *S. sanguinis* T4P, the minor pilins PilA, PilB and PilC^23, 24^. The modular pilins PilB and PilC were proposed to be located at the pilus tip because their bulky adhesin modules are incompatible with polymerisation in the filament body^23, 24^. The non-modular pilin PilA was also predicted to be tip-located since it strongly interacts with (and stabilises) PilC^24^. An AlphaFold^32, 38^-computed PilABC model confirmed that the three minor pilins can coexist in one complex. In this helical complex^24^ – PilA is added first, interacts with PilC, which interacts with PilB – the grafted adhesin modules in PilB and PilC, cap the pilus^23, 24^. We therefore used our filament structure to produce a complete model of *S. sanguinis* T4P by fitting the complex of minor pilins at the tip of the filaments. This was done in two steps.

Since the predicted PilABC architecture implies that PilB connects the complex with the filament, we first positioned PilB in our filament structure. Because PilB and PilE1/PilE2 have canonical SP3 with homologous α1N^22^, we produced a PilB model with a portion of α1N melted (Fig. 7A) using SWISS-MODEL^39^. PilB was then positioned into the filament by aligning its α1-helix with the α1-helix of the S_5_ subunit (counted from the base) rather than the subunit at the apex, to allow the following subunits to serve as reference for aligning the PilAC complex. The alignment of the helices in PilB and PilE1/PilE2 was perfect (Fig. 7B). Next, since the PilAC structural model has been experimentally validated^24^, it was docked as such by aligning the α1-helix of PilC – which also exhibits a canonical SP3^22^ – with the α1-helix of the S_6_ subunit. Strikingly, this resulted in the α1-helix of PilA aligning with the α1-helix of the S_7_ subunit and maintaining the helical symmetry of the pilus, which strongly strengthens the validity of this model (Fig. 7C). Finally, by removing S_5_ and the subsequent major pilin subunits, we produced the final model of a PilABC-capped T4P (Fig. 7D). Critically, although PilA is the first subunit from top, the pilus is capped by the adhesin modules in PilB and PilC, which resemble open wings^24^.

**Fig. 7.**
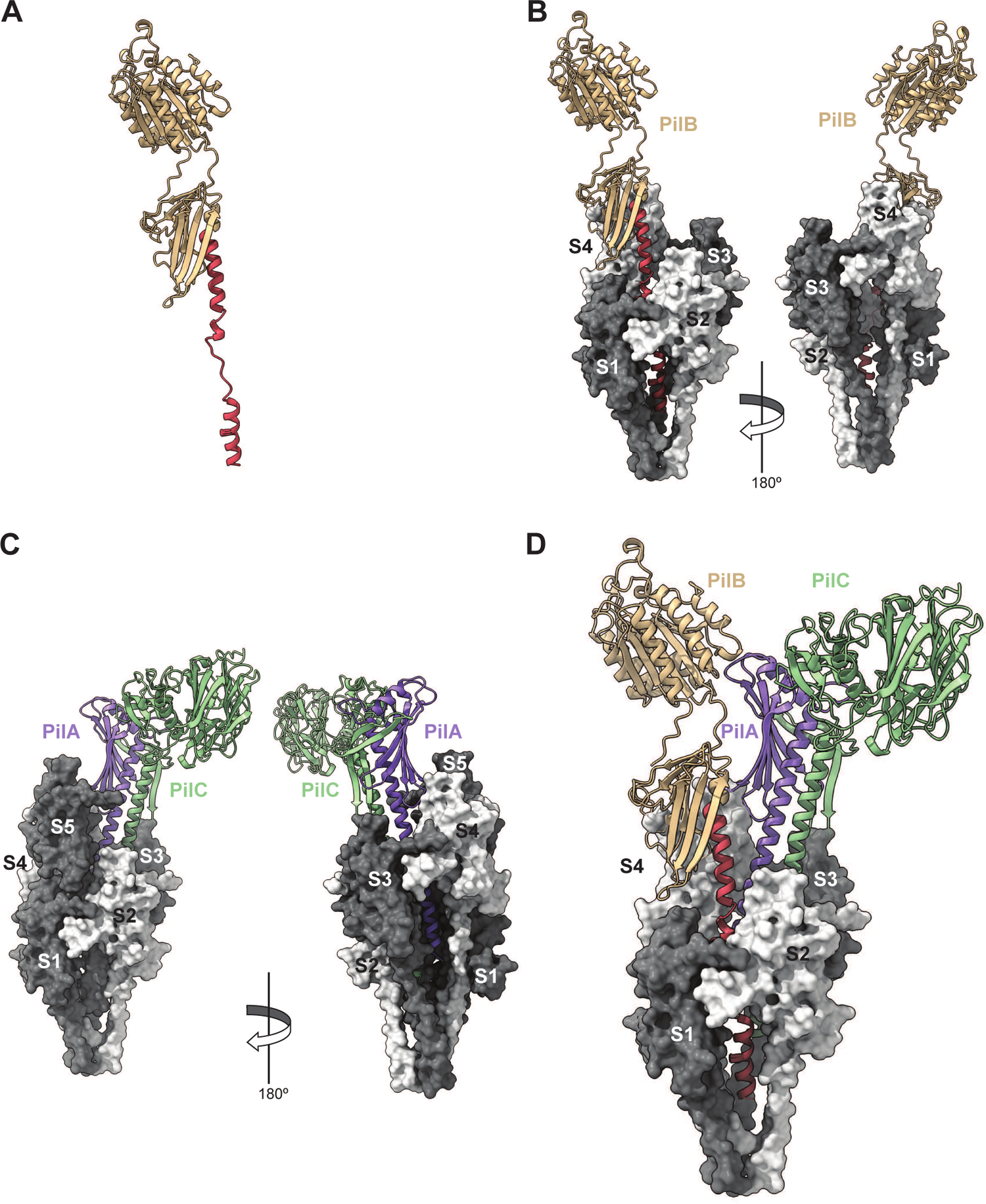
Integrated atomic model of the heteropolymeric T4P in *S. sanguinis* with its tip-located complex of minor pilins. **A**) PilB model with a portion of α1N melted. **B**) PilB superposition with PilE1 subunit S_5_ in the filament structure (180° views). **C**) Superposition of PilAC with PilE1 subunits S_6_ and S_7_ in the filament structure (180° views). **D**) Final model of the T4P pilus in *S. sanguinis*.

By providing a picture of a complete T4F at atomic resolution, including all its pilin subunits, our model of *S. sanguinis* T4P is invaluable. It shows how minor pilins cap the pilus tip efficiently, in this case presenting the adhesin modules in PilB and PilC, which mediate binding of *S. sanguinis* to host cells and structures^23, 24^. This has direct implications for most T4F, which are similarly capped by complexes of minor pilins, playing a variety of roles extending well beyond adhesion.

## Discussion

T4F – a superfamily of filamentous nanomachines ubiquitous in prokaryotes – have been studied for 40 years, primarily in a handful of closely related diderm species. However, mechanistic aspects of their intricate biology remain poorly understood. which led to the development of phylogenetically distant models, recently opening new research avenues. In particular, the monoderm bacterium *S. sanguinis* emerged as a frontline T4F model^19^ because all its major and minor pilin subunits have been structurally and functionally characterised^22–24^. In this study, we have used cryo-EM to determine the structure of *S. sanguinis* T4P, which shines new light on T4F by leading to the notable findings discussed below.

An important achievement in this study is that we have determined the first T4F structure from a monoderm species, which is also the only known heteropolymeric T4F structure in bacteria. This revealed that *S. sanguinis* T4P share the structural characteristics that define T4F, which is not unexpected, but display unique and intriguing features that are likely to play important functional roles. Like in all other T4F, pilin subunits in *S. sanguinis* T4P are arranged in a helical array, in which the extended NT α1-helices form the filament core, while the opposite face of the globular heads form the filament outer shell. Critically, in each pilin subunit, a portion of the NT α1 helix is melted. This structural feature, which was previously observed in T4F from diderm species^12–17^ that are phylogenetically distant from *S. sanguinis*, is therefore likely to be universal in Bacteria. Interestingly, in *S. sanguinis* T4P, flexibility in the pilin subunits appears to be restricted to the melted region, which might play a direct role in the inherent flexibility of the filaments. Overall, although the bacterial T4F architecture is conserved, there are significant structural differences among pilins in the number of β-strands in the central β-sheet and size/structure of the flanking αβ-loop and CT region (Fig. 8). This leads to differences in pilin shapes and sizes, which results in diverse width and helicity parameters of the T4F (Fig. 8). Interestingly, while in diderms the pilin globular heads are tightly compacted within the filaments, in *S. sanguinis* T4P they are more loosely connected with gaps between them. It is possible that this property too contributes to the intrinsic flexibility of *S. sanguinis* T4P. Such flexibility is expected to have an impact on the T4P-mediated properties– twitching motility^20^ and adhesion to host cells^23, 24^ – by facilitating the movement of bacteria and enhancing interaction with surfaces and/or host cells.

**Fig. 8.**
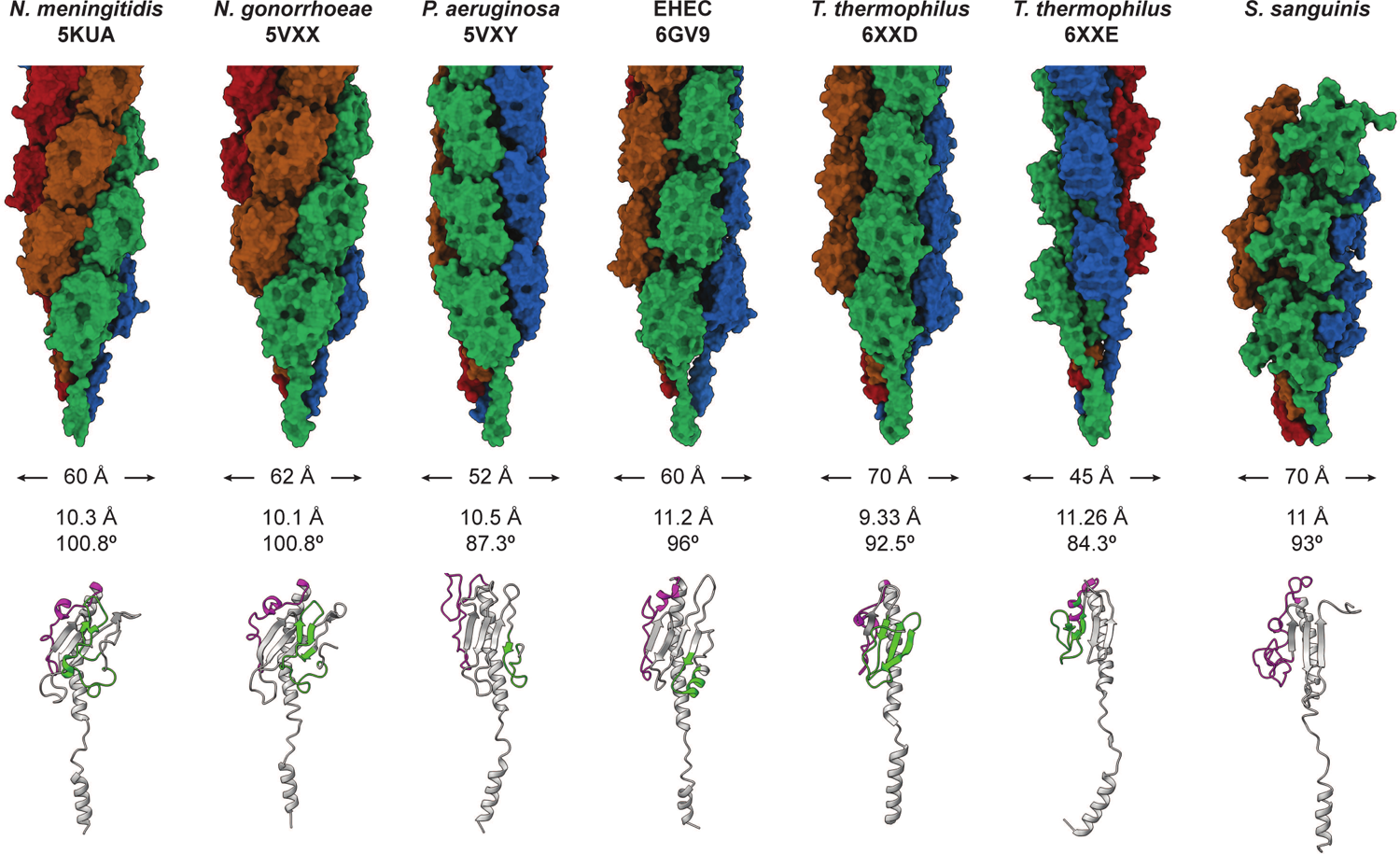
Comparison of known bacterial T4P structures. From Left to right, *N. meningitidis* ^1^, *N. gonorrhoeae*^2^, *P. aeruginosa*^2^, *EHEC*^3^, *T. thermophilus*^4^ (PilA4 and PilA5 T4P) and *S. sanguinis* (present work). **Top**, pilus structure with the pilins coloured along the 4-start helix. The PDB identifiers, diameter and helical symmetry operators are indicated for each filament. **Bottom**, structure of pilin subunits. The pilins have been extracted from the PDB structure of the corresponding filament. For each pilin, the αß-loop is coloured in magenta and the D-region (if present) in green.

An important consequence of the above structural organisation is that the surface of T4F is predominantly constituted by the two edges that flank the central β-sheet and differ the most between pilins, namely the αβ-loop and CT region (Fig. 8). T4F surfaces are often further diversified by post-translational modifications (PTM)^40^. This variability is thought to have important consequences on T4F-mediated functions and is likely used to evade the host immune response^41^ or predation by phages^42^. Although *S. sanguinis* T4P share this structural organisation, they exhibit significant differences. The first difference concerns the unstructured 10 aa-long CT tail in *S. sanguinis* major pilins. In monomers, as previously shown by NMR^22^, this tail exhibits a variety of conformations and orientations within the NMR ensemble and is therefore highly flexible, while the rest of the protein is not. This was a puzzling observation since in diderms T4F piliation occurs only when this CT region is stabilised, when “stapled” to the last β-strand in the central β-sheet, either by a disulfide bond (hence its well-known “D-region” moniker) or by coordination of a metal^5^. Instead, our structure of *S. sanguinis* T4P reveals that the CT tail of pilin subunits is stabilised via a completely different mechanism, and only upon polymerisation within a filament. In brief, the CT tail in one subunit attaches to the αβ-loop of the next through a velcro mechanism, involving a series of electrostatic interactions. The second difference is that unlike in diderms, there are no PTM on the edges in *S. sanguinis* major pilins^22^, but the surface the pilus is diversified in radically different fashion, *i.e*., by a stochastic polymerisation of the two major pilins in the pilus. This ensures that the pilus surface is highly mosaic, which should promote better immune evasion than a highly regular arrangement of the two major pilins. However, a heteropolymeric pilus could confer additional evolutionary advantages, e.g., by modulating filament stability and/or T4P-mediated functions, as proposed for the heteropolymeric T4F in *M. villosus*^28^. Accordingly, in *S. sanguinis*, single Δ*pilE1* and Δ*pilE2* mutants are piliated (piliation is abolished only in the double Δ*pilE1*Δ*pilE2* mutant), but they produce less filaments than the parental strain^20^. Moreover, although they exhibit twitching motility these single mutants move at different speeds than the wild-type strain^22^.

The fact that all known T4F are composed of major and minor pilins^2^ makes our complete structural model of *S. sanguinis* T4P – encompassing all the minor pilins that are key players in T4F biology – the second major achievement in this study. We were in a privileged position to achieve this because, unlike in most other systems, all the minor pilins of *S. sanguinis* T4P (PilA, PilB, and PilC) have been previously characterised structurally and functionally^22–24^. Our atomic model has implications for most (if not all) T4F, but there is an important caveat: it is not an experimentally determined structure, and it is thus perfectible. For example, the adhesin modules in PilB and PilC are almost certainly highly flexible because of the unstructured loops that connect them to their pilin modules. Therefore, the PilB and PilC wings will be “flapping”, which is expected to maximise bacterial adhesion. Second, although it is data-driven, the proposed architecture of the pilus tip and the PilABC complex remains an educated guess. Indeed, because its SP3 is similar to PilE1, we opted to model PilB with a melted α1N portion. In contrast, because the interaction interface between the globular domains of PilA and PilC has been experimentally validated by NMR^24^, we decided to dock the PilAC AlphaFold model as such. However, since the SP3 of PilC is similar to PilE1 it cannot be excluded that the α1N of PilC too is melted upon filament polymerisation, which is not likely for PilA that has a highly peculiar SP3^22^. Determining an atomic resolution structure of a T4F with its tip-located minor pilins would shed light on these issues, but this would require methodological and technical advances beyond the state of the art. These advances include pilus preparation methods that would preserve the integrity of the filaments from base to tip, or electron cryotomography improvements allowing the structural characterisation of filaments emanating from cells at near-atomic resolution.

The complete model of *S. sanguinis* T4P, including three minor pilins at the tip, has several important implications. The filament starts with PilA, the only pilin that lacks a Glu_5_. This is consistent with the structural characterisation of a complex of four minor pilins widespread in T4F^43–45^, generically known as HIJK^46^, which is capped by the K subunit that also lacks Glu_5_. Since the negatively charged Glu_5_ in subunit S usually establishes an important salt bridge with the positively charged N-terminus of the S_+1_ subunit, it is not unexpected that this residue is not needed (and therefore absent) in the subunit at the apex of the pilus, because there is no subunit in +1 position. Since PilA specifically interacts and stabilises PilC^24^, this is the next pilin in the filament. PilC has an unusually large modular pilin with a lectin domain, which bind glycans prevalent in the human glycome^24^. The AlphaFold model of PilAC, which confirmed the interaction interface characterised experimentally^24^, fitted very well in the filament structure. PilA is thus an anchor for PilC, facilitating the presentation of this adhesin at the tip of filaments. The structural homology between PilA and the I subunit of the HIJK complex^24^ suggests that this latter complex might play a similar role – facilitating the presentation at the tip of effectors such as PilC/PilY1 in T4P from diderms^47^, or secreted effectors in T2SS^48^. The third pilin in the filament is PilB, another large modular pilin and *bona fide* adhesin with a vWA module that binds protein ligands such as fibronectin and fibrinogen^23^. The AlphaFold model of PilB, in which we melted a portion of α1N, fitted very well on its own in the filament structure. This led to significant remodelling of the interface between PilB and PilC proposed by AlphaFold in the PilABC complex^24^, which interestingly could not be fitted as such. This is reminiscent of the melting in α1N of pilins occurring during filament assembly, which similarly could not be predicted but was demonstrated by cryo-EM in multiple T4F^12–17^, including in *S. sanguinis* in this study.

In conclusion, by solving the structure of a T4F in a bacterial species radically different from previously characterised ones and producing a complete picture including the minor pilins (key but often poorly characterised players in T4F biology), this study has general implications for T4F. Moreover, it definitely cements *S. sanguinis* as a T4F model and paves the way for further investigations that will improve our understanding of these fascinating filaments.

## Material and methods

### Strains and growth conditions

*E. coli* strains were grown in liquid or solid lysogeny broth (LB) medium (Difco), with spectinomycin (100 µg/ml), kanamycin (50 µg/ml) and/or ampicillin (100 µg/ml), when required. All antibiotics were from Sigma. Strain DH5α was used for cloning, while strain BTH101 (Euromedex) – a non-reverting *cya* mutant – was used in BACTH assays^29^. We amplified the full-length genes *pilE1* (5’-cgc<u>ggatcc</u>CATGTTAAACAAATTACAAAAATTCCG-3’ and 5’-cgc<u>ggtacc</u>GCGTTTGAGTTTACACCATTAGCA-3’) and *pilE2* (5’-cgc<u>ggatcc</u>CATGTTAAACAAATTGCAAAAATTCCG-3’ and 5’-cgc<u>ggtacc</u>GCTTTTGAATTAGCACCAGCTTC-3’) from genomic DNA of *S. sanguinis* 2908^20^ using high-fidelity Pfu DNA polymerase (Agilent). PCR products were cloned directly into pCR8/GW/TOPO (Invitrogen). Inserts were verified by sequencing, cut out from the TOPO derivative by *Bam*HI and *Kpn*I digestion, gel-extracted, and subcloned into the BACTH vectors pUT18C and pKT25. Cloning was carried out using standard molecular biology techniques^49^.

*S. sanguinis* was grown as described^20^ on plates containing Todd Hewitt (TH) broth (Difco) and 1 % agar (Difco), or in liquid culture in THTH, *i.e.*, TH broth containing 0.05 % tween 80 (Merck) to limit bacterial clumping, and 100 mM HEPES (Euromedex) to prevent acidification of the medium. When required, 500 μg/ml kanamycin was used for selection. Plates were incubated at 37°C in anaerobic jars (Oxoid) under anaerobic conditions generated using Anaerogen sachets (Oxoid), while liquid cultures were grown statically under aerobic conditions. To construct the *S. sanguinis* Δ*fim* mutant, we deleted the locus involved in the production of sortase-assembled pili^25^. We replaced the genes from SSV_1503 to SSV_1499 (*srtC*) by a promoterless *aphA-3* cassette, which confers resistance to kanamycin. To do this, we fused by splicing PCR the regions upstream (5’-GCCAAGCACCTGACTAGTAG-3’ and 5’-ggtgatattctcattttagccatTATAATCTCCTAATTTTATCTTCACTC-3’) and downstream the *fim* locus (5’-ttttactggatgaattgttttagGGAAAAGAAAAGAGCCGAGC-3’ and 5’-ATTCCACCGCGTCATCAATG-3’) to *aphA-3* (5’-ATGGCTAAAATGAGAATATCACC-3’ and 5’-CTAAAACAATTCATCCAGTAAAA-3’). We directly transformed the PCR product into strain 2908 and selected allelic exchange mutants on kanamycin plates. Allelic exchange was confirmed by PCR.

#### T4P visualisation

Surface-associated T4P in *S. sanguinis* Δ*fim* were visualised by TEM after negative staining as follows. Bacteria were grown in THTH until OD_600_ 0.8, adsorbed for 3 min to glow-discharged carbon-coated grids (EMS), and fixed 5 min in 2 % glutaraldehyde. The grids were cleaned by floating them sequentially 10 times on drops of pilus buffer (20 mM Tris, pH 7.5, 50 mM NaCl), and then stained for 2 min with 2 % aqueous uranyl acetate. Stain solution was gently drained off the grids, which were air-dried before visualisation using a Tecnai 200KV electron microscope (Thermo Fisher Scientific). Digital image acquisition was made with a numeric camera (16 megapixel, CMOS, Oneview, Gatan).

#### T4P purification

T4P were purified from *S. sanguinis* as described elsewhere^20^ with minor modifications. Liquid cultures grown O/N in THTH were used to re-inoculate pre-warmed THTH and grown statically until the OD_600_ reached 1. Bacteria were pelleted by centrifugation for 10 min at 4,149Lg at 4°C. Pellets were re-suspended in ice-cold pilus buffer by vigorous pipetting up and down, which was enough to shear T4P. Bacteria were then pelleted as above, and supernatant containing the pili was transferred to a new tube. This centrifugation step was repeated, before the supernatant was passed through a 0.22 µm pore size syringe filter (Millipore). Pili were then pelleted by ultracentrifugation, resuspended in pilus buffer, and tested by SDS-PAGE/Coomassie, essentially as described^20^.

#### Cryo-EM sample preparation and data acquisition

R2/2 Cu 200 mesh grids (Quantifoil) were glow-discharged for 40 s at 2.7 mA. Then, 4 µl of freshly purified pili were applied and the excess of sample was immediately blotted away (3.5 s blot time, 4°C chamber temperature, 100% humidity) in a Vitrobot Mark IV (Thermo Fischer Scientific), before being plunge-frozen in liquid ethane. Cryo images of the purified pili were recorded with a Talos Arctica microscope (Thermo Fischer Scientific) operated at 200 kV and equipped with a K2 summit direct electron detector (Gatan). Dose fractioned data were collected in a defocus range of −0.4 to −1.4 µm at 45,000 x magnification, corresponding to a pixel size of 0.93 Å using SerialEM^50^. The total dose was 50.74 electrons/Å^2^, and the dose per frame was 1.34 electrons/Å^2^.

#### Image processing

All data processing was carried out in Cryosparc^51^. Movies were aligned for beam-induced motion using the function “Patch motion correction”, while CTF (Contrast Transfer Function) parameters were assessed using “Patch CTF Estimation”. Non-overlapping segments of the *S. sanguinis* T4P were manually picked, and particles were extracted using a box size of 320 pixels. These particles were 2D classified, the best 2D classes were selected and used as references to automatically pick the filament in all the micrographs. This was done using the program “Filament Tracer”^52^ by indicating a filament diameter of 70 Å, a separate distance between segments of 0.7 and a minimum filament length to consider of 1. After extraction, several rounds of 2D classification were performed and 393,556 particles, corresponding to well-resolved classes were selected for further processing. Four *ab initio* models were generated and a single best one was used as 3D reference for a homogenous refinement. A mask was created using the function “Volume Tool” for the central part of the filament, which was used for several rounds of local refinement. This processing led to a final map at 3.67 Å of resolution.

#### Building and refinement of atomic models

Model of the major pilin PilE1 was generated using Alphafold^32^ and docked into the refined cryo-EM map using the program “Fit in map” from the software ChimeraX^34^. The map was sharpened in PHENIX^31^ (phenix.autosharpen)^53^, and the final model was refined by several rounds of manual refinement in ISOLDE^33^ and real-space refinement using phenix.real_space_refine^36^. The model was validated using MolProbity^54^ and phenix.validation_cryoem^55^ implemented in the PHENIX software.

#### Testing interaction between PilE1 and PilE2 by BACTH

BACTH assays as described elsewhere^30^ with minor modifications. In brief, BTH101 cells, co-transformed with pairs of recombinant pUT18C and pKT25 plasmids, were plated on selective MacConkey plates supplemented with 0.5 mM IPTG and 1 % maltose (Sigma). Plates were incubated at 30°C and the colour of the colonies was scored after 40 h. Each assay was repeated three times with a positive and a negative control included. The efficiency of the functional complementation between T18 and T25 was quantified by measuring ß-galactosidase activity in liquid culture as previously described^30^. Single colonies were picked from the above MacConkey plates after 48 h of growth, inoculated in 5 ml LB supplemented with 0.5 mM IPTG and antibiotics, and the bacteria were grown O/N at 30°C. The next day, the cultures were diluted in M63 broth and the OD_600_ was recorded. Cells were permeabilised with chloroform and SDS^30^ during for 40 min at 30°C with shaking at 250 rpm. Ten µl of the permeabilised cells were then diluted into 990 µl PM2 buffer containing 100 mM ß-mercaptoethanol, and incubated at 28°C for 5 min^30^. The ß-galactosidase reaction was started at 28°C with O-nitrophenol-ß-galactoside diluted in PM2 buffer and stopped with 500 µl 1 M Na_2_CO_3_ after 20 min for positive samples, or 60 min for negative samples. The OD_420_ was recorded, and the enzymatic activity A (units/ml) was quantified as A = 200 x (OD_420_/min of incubation) x dilution factor. The results were expressed as enzymatic activity per milligram of bacterial dry weight (U/mg), so that 1 unit corresponds to 1 nmol of ONPG hydrolysed per minute at 28°C^30^. The assay was performed on three independent cultures for each plasmid combination.

#### Bioinformatics and modelling

PilB’s NT α-helix (residues 1-58) is similar in sequence to the corresponding portion in PilE1/E2 (25 % identity). Therefore, to model this part of PilB that is the main assembly interface in the pilus, we performed homology modelling with the NT portion (residues 1-58) of PilE1 using SWISS-MODEL^39^. We then superimposed the NT α-helix in PilB model with the NT α-helix in the S_5_ PilE1 subunit, close to the centre of our filament structure. Similarly, we superimposed the NT α-helix of PilC in the AlphaFold PilAC model^24^ with the NT α-helix in the next PilE1 subunit S_6_, which led to superposition of PilA’s NT α-helix with the NT α-helix of S_7_ PilE1 subunit. The final model was obtained by removing S_5_ and the subsequent major pilin subunits. The quality of the computed model was estimated using the Structure Assessment tool in SWISS-MODEL.

## Acknowledgements

This work was funded by the Agence Nationale de la Recherche (ANR-21-CE11-0008-01 to VP and RF) and the Medical Research Council (MR/P022197/1 to VP). We thank Sophie Helaine (Harvard Medical School) and Romé Voulhoux (Laboratoire de Chimie Bactérienne, Marseille) for critical reading of the manuscript.

## Legends to Supplementary Figures

**Fig. S1. Filaments in purified pilus preparations from *S. sanguinis***. **A**) Representative cryo-EM micrograph of a purified pilus preparation from *S. sanguinis*. The three types of filaments are indicated by arrowheads: white arrowhead (6 nm-wide filaments corresponding to native T4P), red arrowhead (12 nm-wide “thick” filaments corresponding to T4P), and yellow arrowhead (3 nm-wide “thin” filaments corresponding to DNA). **B**) Analysis of the thick filaments. **Top**, averages after 2D classification. **Bottom**, orthogonal views of the cryo-EM map that was generated. **C**) Analysis of the thin filaments. **Top**, averages after 2D classification. **Bottom**, cryo-EM map in which the B-DNA structure was docked.

**Fig. S2. Resolution of the cryo-EM map of *S. sanguinis* T4P upon imposition of helical symmetry. A**) **Left**, representative average of pilus sections from cryo-EM micrographs after 2D classification. **Right**, slice of the mask used for 3D refinement of the central portion. **Bottom**, estimation of the resolution by gold standard FSC. **B**) Density map sharpened with DeepEMhancer^35^. Side and end views of the filament are presented.

**Fig. S3. Sequence conservation between PilE1 and PilE2 in the context of the cryo-EM map.** The conservation between the two pilins was estimated with ConSurf. Orthogonal views of the pilin (**A**) and the (**B**) T4P in surface representation. The regions that are conserved between PilE1 and PilE2 are in purple, whereas the regions that differ are in cyan. The regions with no data are coloured in grey. **C**) The regions that differ between PilE1 and PilE2 correspond to areas of poorer local resolution in the 3.7 Å cryo-EM density map of *S. sanguinis* T4P.

**Fig. S4. Flexibility between the different subunits in the T4P structures.** Superposition of the eight complete pilin subunits which were resolved in the pilus structure.

**Fig. S5. Areas of interaction between the pilin subunits in the structure of *S. sanguinis* T4P. Top,** Surface representation of a pilin subunit located in the central part of the filament shown in two orientations. The interaction areas with the neighbouring subunits are coloured in orange. **Bottom**, interactions between the S subunit and the neighbouring subunits in the 1-start, 3-start and 4-start helices (180° views).

## Notes

### Competing Interest Statement

The authors have declared no competing interest.

