## Supplementary figures and images for "Structure of a heteropolymeric type 4 pilus from a monoderm bacterium"

### Supplementary Figure 1

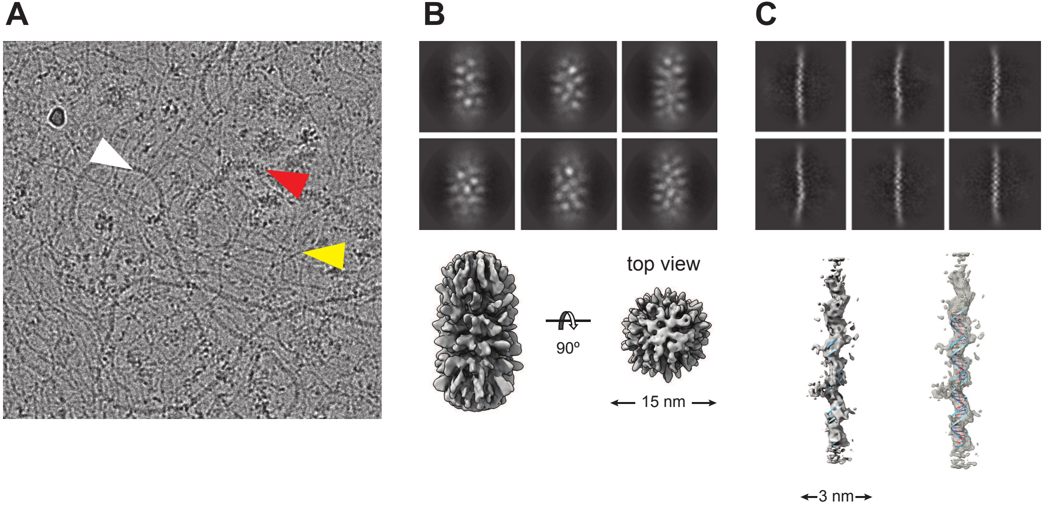

### Supplementary Figure 2

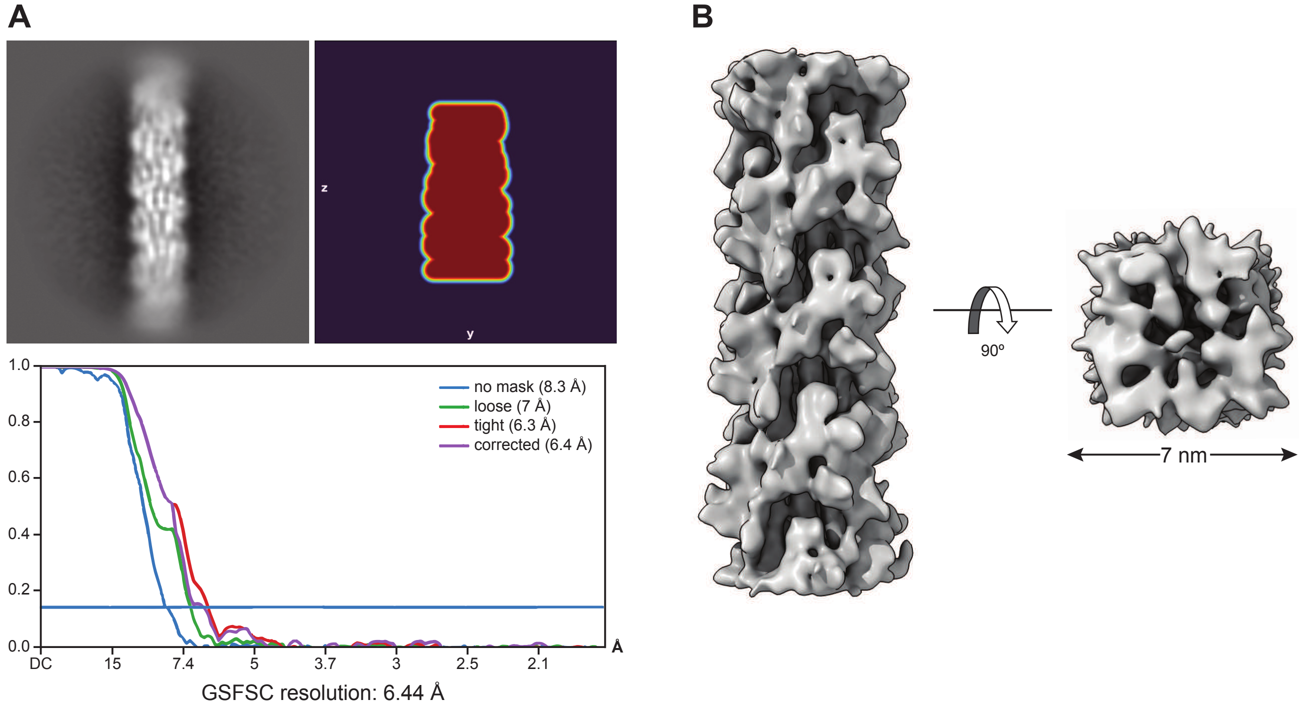

### Supplementary Figure 3

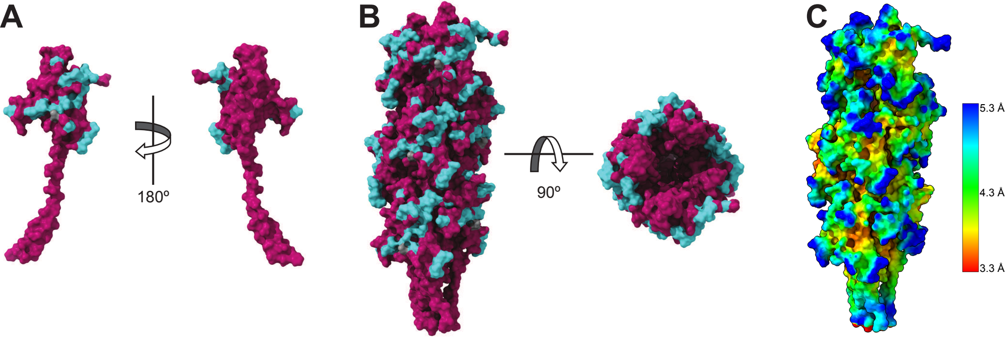

### Supplementary Figure 4

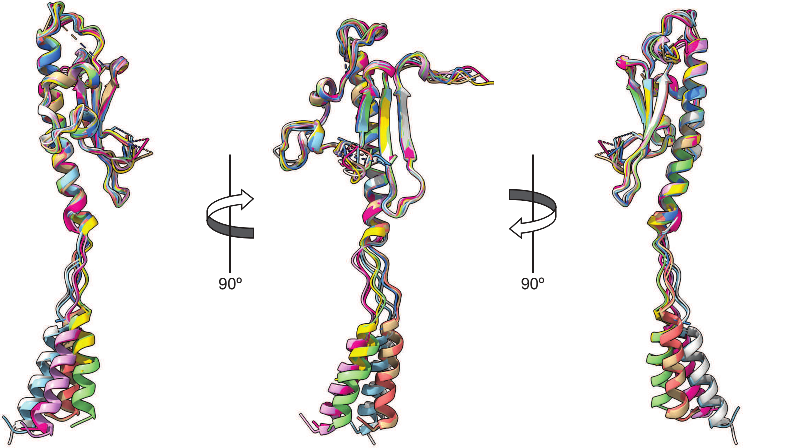

### Supplementary Figure 5

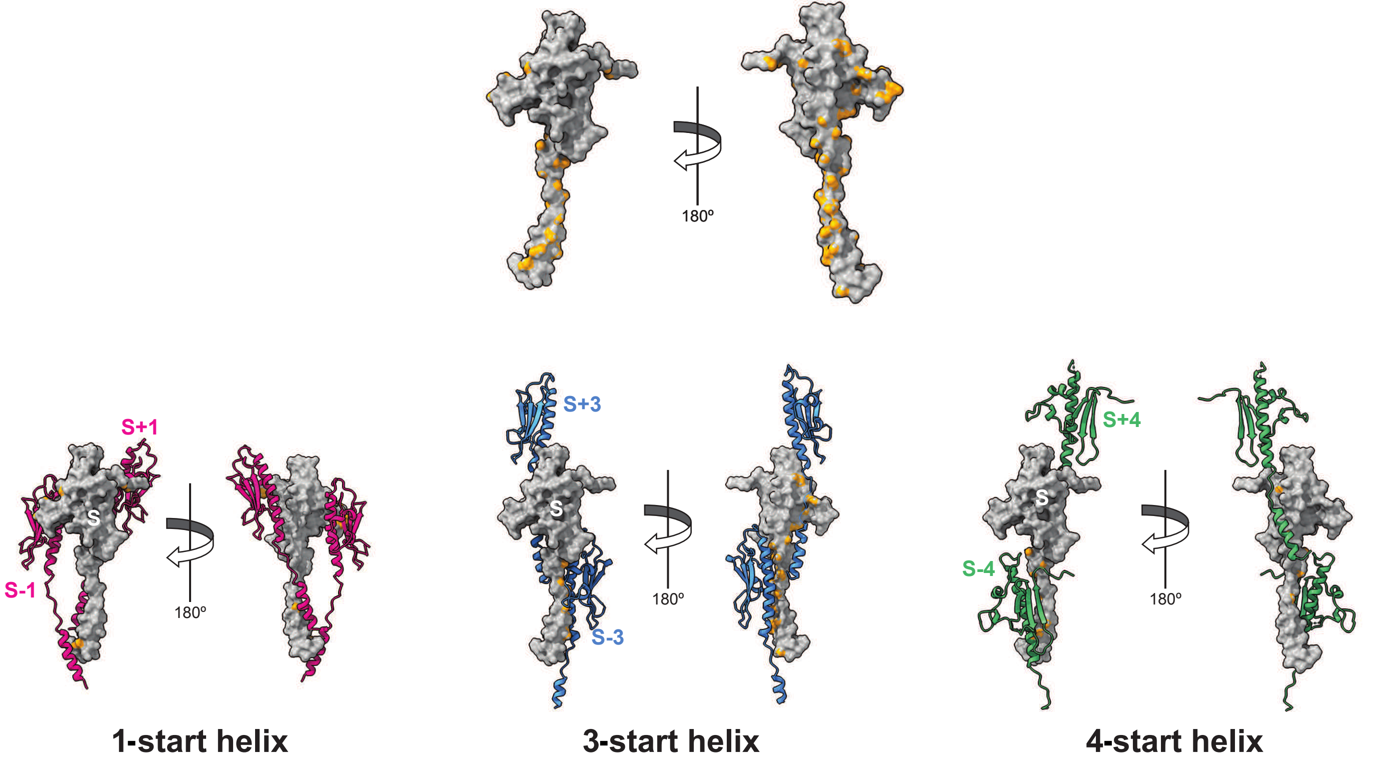
